# Reduced neuronal self-avoidance in mouse starburst amacrine cells with only one *Pcdhg* isoform

**DOI:** 10.1101/2025.05.29.656828

**Authors:** Cathy M. McLeod, Seoyoung Son, Muhammad Nazmul Haque, Andrew M. Garrett

## Abstract

The clustered protocadherins (cPcdhs) are a family of ∼60 homophilic cell adhesion molecules expressed across three gene clusters (*Pcdha, Pcdhb*, and *Pcdhg*) with a variety of essential roles in the developing nervous system. Some of these roles rely on specific isoforms, while others are more consistent with a model of isoform redundancy or a requirement for diversity. The γ-Pcdhs (expressed from the *Pcdhg* gene cluster) are particularly important for neuronal self-avoidance in starburst amacrine cells in the mouse retina. Here, we used mouse mutants to test two of the C-type isoforms – γC4 and γC5 – and found that neither was required for normal self-avoidance. Conversely, when we analyzed a mutant with only γC4 intact, we found significant failures in self-avoidance that could not be completely rescued by overexpression of this isoform from a transgene. We have recently found that this isoform is essential for normal neuronal survival during development, and our new findings here support the hypothesis that γC4 is specialized for the survival function at the expense of a significant role in self-avoidance.

## INTRODUCTION

Self-avoidance is an essential mechanism in neuronal morphogenesis. As developing dendrites and axons branch and elaborate, self-avoidance promotes neurites from the same cell (“self”) or same cell type (“homotypic”) to move away from each other and sample potential “non-self” interactors rather than becoming entangled with each other. It has been described in both vertebrate and invertebrate systems with a growing list of cell-surface molecules implicated in mediating self-contacts leading to avoidance [1]. Closely related to self-avoidance is the concept of “self/non-self discrimination”, in which individual neurons have a unique recognition signal differentiating “self” processes from any other “non-self” protrusion. This type of individual barcoding requires an extremely diverse signal exemplified by two families of cell adhesion molecules (CAMs): the 38,000 isoforms generated via alternative splicing of the *dscam1* gene in *Drosophila* and the ∼60 isoforms generated by the clustered Protocadherin (cPcdh) locus in vertebrates [2].

In mouse, the cPcdhs comprise three gene clusters organized across nearly 1 Mb on chromosome 18: *Pcdha* encodes for 14 α-Pcdhs, *Pcdhb* for 22 β-Pcdhs, and *Pcdhg* for 22 γ-Pcdhs [3, 4]. *Pcdha* and *Pcdhg* have a similar structure with individual variable exons encoding the entire extracellular domain, transmembrane domain, and variable cytoplasmic domain of individual isoforms, while three cluster-specific constant exons encode a common C-terminal cytoplasmic domain. β-Pcdhs are encoded by single exons for each isoform with no constant domain. Isoforms are further subdivided based on sequence homology: There are five C-type isoforms – αC1 and αC2 encoded in the *Pcdha* cluster and γC3, γC4, and γC5 by the *Pcdhg* cluster. The other γ-Pcdhs isoforms are categorized as γA or γB subtypes. All isoforms are CAMs in the cadherin superfamily with homophilic binding specificity in *trans* [5, 6]. They have less isoform specificity when dimerizing in *cis* resulting in lattices of dimers between membranes to confer recognition specificity in a combinatorial manner [7-11]. Thus, when expressed in stochastic isoform combinations, they can generate recognition signals with extreme diversity.

cPcdhs are not only engaged in self-avoidance, but they also contribute to neural developmental functions including dendrite arborization [12, 13], neuronal survival [14-16], and synaptic development [17-19]. In recent years it has emerged that some of these require specific isoforms, particularly the C-type isoforms, rather than cluster diversity or redundancy. For example, we previously found that γC3 is crucial for dendrite arborization in the cerebral cortex [20, 21] and that γC4 is the only isoform essential for normal neuronal survival [22-24]. Even in the case of self-avoidance single isoforms can be dominant, as in the projections of serotonergic axons which rely on αC2 for avoidance leading to dispersion [25, 26]. γ-Pcdhs are particularly important for self-avoidance in starburst amacrine cells (SACs) in the retina and Purkinje neurons in the cerebellum [27]. In SACs, loss of expression from *Pcdhg* but not *Pcdha* resulted in loss of normal self-avoidance, while disruption of both clusters caused a more severe phenotype than loss of *Pcdhg* alone [28]. However loss of the three γC type isoforms in an allele compound heterozygous with a conditional allele disrupting the whole cluster did not result in self-avoidance failures, and expression of a single isoform from a transgene (γA1 or γC3) was able to rescue loss of the whole cluster [27, 29]. Together, this suggests a model of redundancy for the γ-Pcdh isoforms in self-avoidance *per se*. Consistent with this, we previously studied a mutant for *Pcdhgc3* and did not find indication of self-avoidance failures in SACs [21]. Here, we investigated *Pcdhgc4* and *Pcdhgc5* mutants and found that neither was essential for SAC self-avoidance. Conversely, when we analyzed mutants with only *Pcdhgc4* left intact but lacking the other 21 isoforms, we found significant reductions in self-avoidance that could not be completely rescued by overexpression of transgenic *Pcdhgc4*.

## RESULTS

### Individual C-type isoforms are not required for ON SAC self-avoidance

To test if γC4 or γC5 are essential contributors to ON SAC self-avoidance, we analyzed retinas in two mouse mutants. *Pcdhg*^*C5KO*^ is a recently described allele harboring a 7 base-pair deletion in the *Pcdhgc5* exon [30]. These mice are overtly normal, have a normal lifespan, and can be analyzed as homozygous *Pcdhg*^*C5KO/C5KO*^ mutants. Due to the essential role that γC4 serves in promoting neuronal survival, *Pcdhg*^*C4KO/C4KO*^ homozygous mutants only survive for a few hours after birth [22, 23]. Therefore, to analyze later developing morphologies like those of SACs, we made these compound heterozygous with the conditional allele *Pcdhg*^*fcon3*^ to make *Pcdhg*^*C4KO/fcon3*^ [14, 16]. SACs are the only cholinergic neurons in the retina, so to limit recombination of the conditional allele to SACs, we crossed these mice with *Chat*^*Cre*^, which harbors an IRES-Cre cassette knocked into the endogenous *Chat* locus [31].

*Chat*^*Cre*^ was also useful for labeling SACs in conjunction with Brainbow AAV vectors. These are fluorescent Cre reporters that express a stochastically chosen color after recombination [32]. We injected a 1:1 mixture of the two available vectors to label in four colors (two per vector) into the vitreous of *Chat*^*Cre*^ eyes. After two weeks, we isolated the retinas, immunostained for the antigenically distinct fluorescent proteins, and imaged SACs in whole mount retinal preparations. This multicolor approach allowed us to achieve sparse labeling to image individual SACs in isolation. We analyzed SACs for self-avoidance defects with two methods: a unitless radial asymmetry index, and a qualitative Elo scoring method which leverages the power of the human eye to capture differences in morphology that are difficult to measure such as more discrete fasciculation Briefly, the radial asymmetry index is a measure of how SAC dendrites are arranged in 360 degrees around the soma, where a number close to zero indicates there is low variance/ high radial symmetry of the SAC, and higher numbers indicate a higher variability and asymmetry [28, 33]. The Elo score was obtained by having observers, blind to genotype, choose between two randomly presented images by judging which is less fasciculated. Based on “wins” and “losses” in iterative pairwise matchups, the images received scores using an Elo algorithm. Scores above zero (more “wins”) indicate that a SAC has normal self-avoidance, while a score below zero (more “losses”) indicate failure in self-avoidance compared with other SACs [34]. We compared morphologies of ON SACs across three genotypes: *Chat*^*Cre*^::*Pcdhg*^*+/+*^, *Chat*^*Cre*^::*Pcdhg*^*C4KO/fcon3*^, and *Chat*^*Cre*^::*Pcdhg*^*C5K0/C5KO*^. SACs from the two mutant genotypes were indistinguishable from *Pcdhg*^*+/+*^ controls (**Figure 1**).

**Figure 1.**
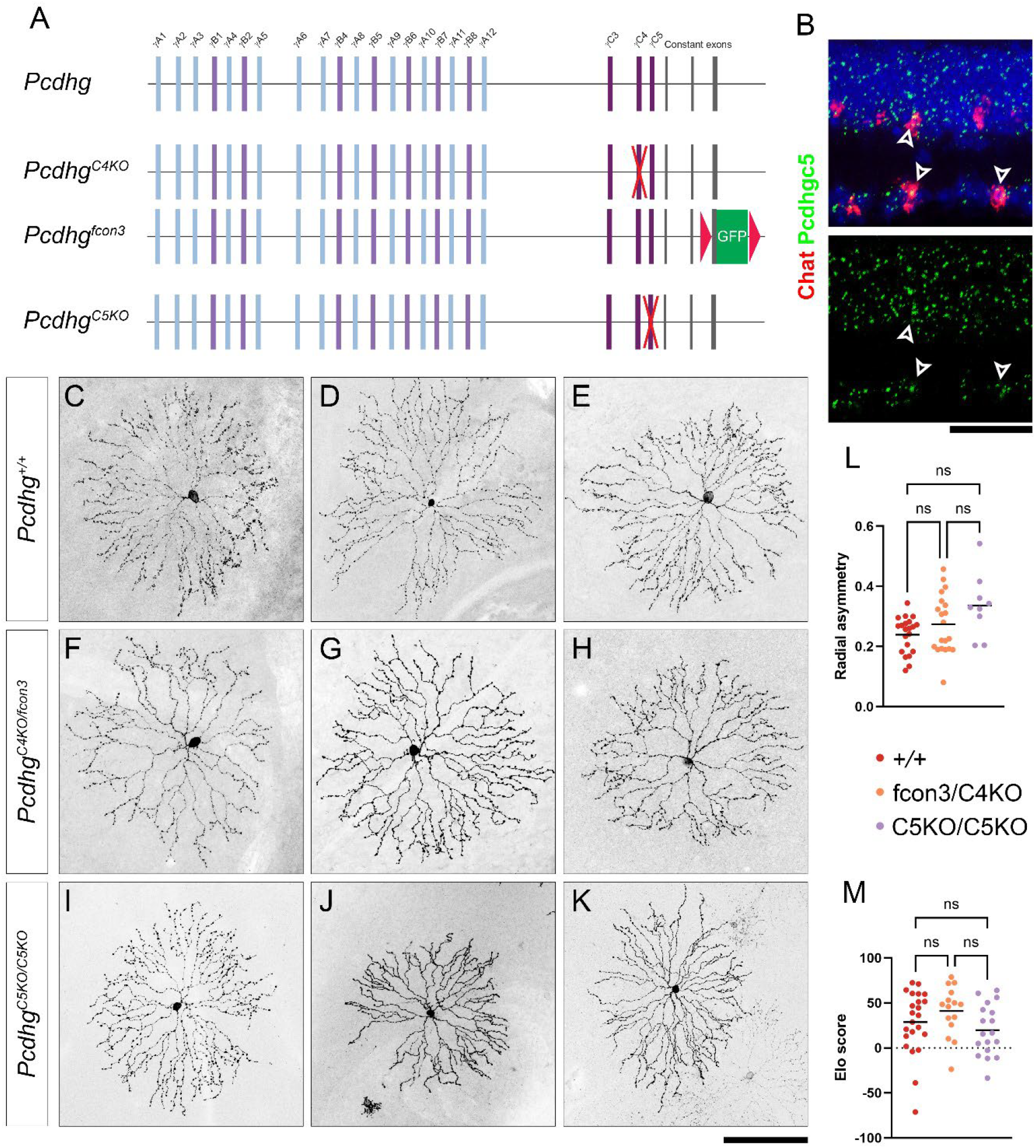
Individual γC-type isoforms are not required for ON-SAC self-avoidance. **A**) A schematic of the *Pcdhg* locus with γA and γB type variable exons at the 5’ end and γC type isoforms at the 3’ end adjacent to the three constant exons. The *Pcdhg*^*C4KO*^ allele harbors a frameshifting 13-bp deletion disrupting the γC4 isoform. To circumvent the neonatal lethality of homozygous mutants, *Pcdhg*^*C4KO*^ is made compound heterozygous with *Pcdhg*^*fcon3*^ and crossed with *Chat*^*Cre*^ to limit the loss of function to starburst amacrine cells (SACs). *Pcdhg*^*C5KO*^ includes a 7-bp frameshifting deletion in the *Pcdhgc5* exon. **B**) RNAscope in adult retina verified that, along with many other retinal cell types, SACs expressed *Pcdhgc5*. Projections of ON SACs labeled with Brainbow in *Chat*^*Cre*^ from (**C**-**E**) *Pcdhg*^*+/+*^, (**F**-**H**) *Pcdhg*^*C4KO/fcon3*^, and (**I**-**K**) *Pcdhg*^*C5KO/C5KO*^ retinas show a range of normal, radially symmetrical morphologies. Individual SAC morphologies were quantified by (**L**) radial asymmetry and (**M**) Elo score. Scale bar is 50 μm in **B**, 100 μm in **C**-**K**.

### Disrupted self-avoidance in Pcdhg^1R1/1R1^ mutant retinas

These results are consistent with the “redundancy” hypothesis that no isoform is essential for SAC self-avoidance and any isoform is sufficient for this process [27]. However, γC4 has several features that make it unique. It alone among γ-Pcdhs cannot form homodimers in *cis* [7], and it is the sole isoform responsible for promoting neuronal survival, which likely involves distinct protein interactions [22, 24]. To test if γC4 is also capable of promoting self-avoidance, we analyzed SACs from *Pcdhg*^*1R1/1R1*^ mutants. This allele includes a large deletion disrupting *Pcdhga1* to *Pcdhgc3* but sparing *Pcdhgc4*, then a frameshifting indel in Pcdhgc5; γC4 is the only γ-Pcdh produced in these mutants (**Figure 2A**) [22]. We crossed these mice with *Chat*^*Cre*^ animals and analyzed individual SACs as with the mutants described above. Unlike in *Pcdhg*^*C5K0/C5KO*^ or *Pcdhg*^*C4KO/fcon3*^ mutants, we found significant failures of self-avoidance in *Pcdhg*^*1R1/1R1*^ SACs as measured both by radial asymmetry and Elo score (**Figure 2J-N**). Similarly to that reported for whole cluster mutants [27], there was variability in the severity of these defects. To assess these defects at the population level, we stained whole mount retinas for Chat, imaged by confocal microscopy, and made maximum projections through the ON plexus but excluding cell bodies. Here, failure in self-avoidance could be observed as interdendritic gaps vacated by SAC neurites as they became entangled with themselves and with each other. We quantified this effect by measuring the maximum interdendritic gap in each image and by using the pairwise comparison approach to generate Elo scores. By both measurements, *Pcdhg*^*1R1/1R1*^ SACs were significantly more fasciculated than controls (**Figure 2B-I**).

**Figure 2.**
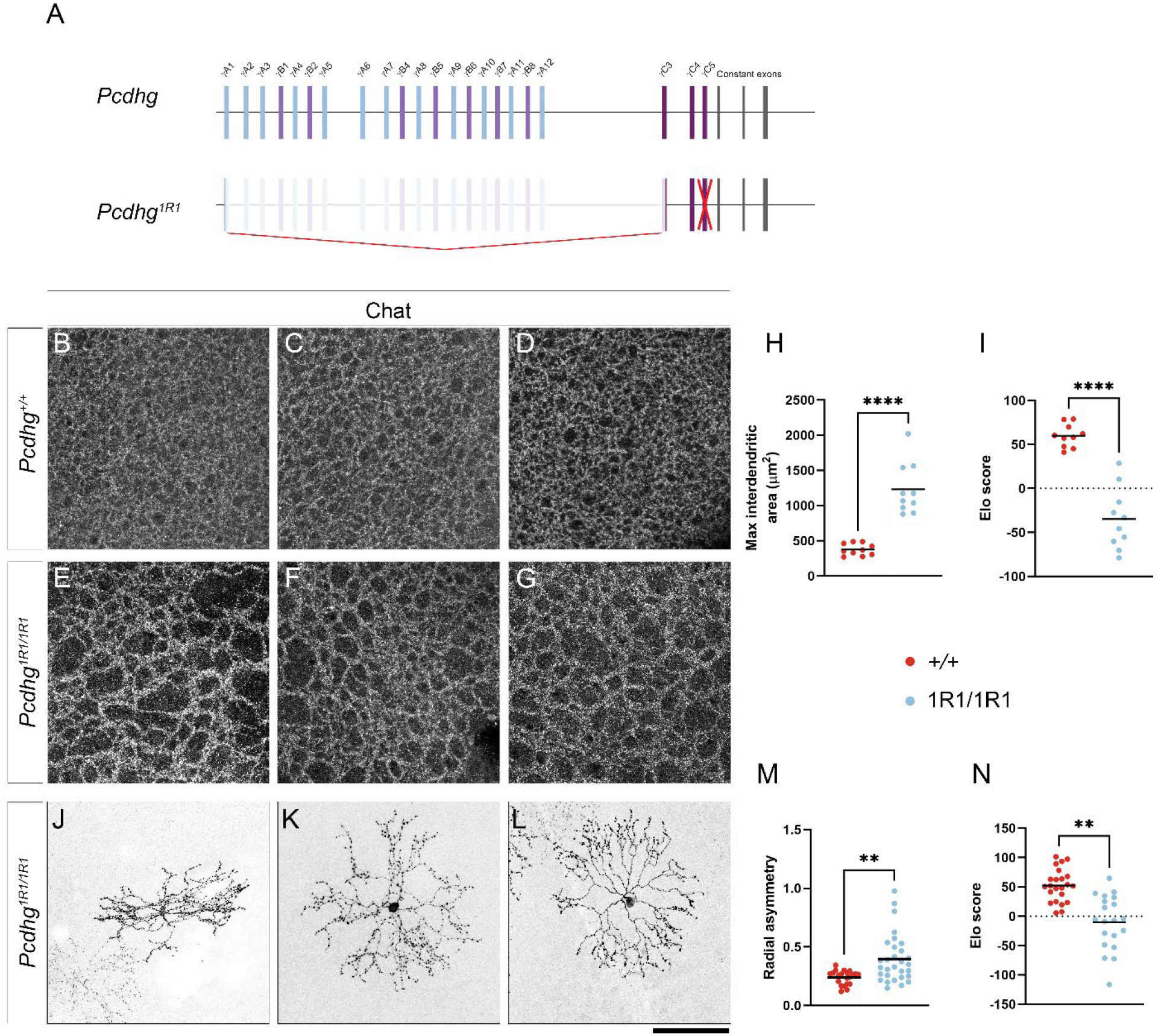
Disrupted self-avoidance in *Pcdhg*^*1R1/1R1*^ mutant retinas. **A**) A schematic representation of the *Pcdhg*^*1R1*^ allele indicates a large deletion from *Pcdhga1* to *Pcdhgc3* and a frameshifting mutation in *Pcdhgc5*, leaving only *Pcdhgc4* intact. Projections through the ON SAC plexus stained for Chat in (**B-D**) *Pcdhg*^*+/+*^ and (**E-G**) *Pcdhg*^*1R1/1R1*^ retinas. Plexus in *Pcdhg*^*1R1/1R1*^ is characterized by large gaps between dendritic fascicles, as quantified in **H** and **I. J-L**) Individual *Pcdhg*^*1R1/1R1*^ SACs labeled with Brainbow in *Chat*^*Cre*^ show self-avoidance failures with a variety of severity, quantified in **M** and **N**. Scale bar is 100 μm. **** is P < 0.0001, ** is P < 0.01 by Tukey (**H,M**) or Dunn’s (**I,N**) multiple comparison test.

This suggests a hypothesis that γC4 alone is not sufficient to promote self-avoidance. However, one caveat is that the total level of γ-Pcdh constant domain is majorly reduced in these mice, and even γC4 protein levels are lower than in wild type animals [22]. Thus, even if γC4 could promote self-avoidance, there may not be enough present to form the complexes required for the dynamic process of self-avoidance during development [33]. To test this alternate hypothesis, we used a recently developed transgenic line, *C4-GFP*, that expresses γC4 with a C-terminal GFP fusion from a CAG promoter after Cre-mediate excision of a stop cassette (**Figure 3A**) [23]. We found that this transgene produces approximately 10-fold expression at the mRNA level and, when expressed in neurons, was able to rescue neonatal lethality in *Pcdhg*^*C4K0/C4KO*^ mice [23]. Of particular importance here, the design of this allele used the same promoter and insertion site (at the *Rosa* locus) as the C3-mCherry and A1-mCherry transgenic lines previously found to rescue self-avoidance after disruption of the entire *Pcdhg* locus [27, 29]. To express the transgene in SACs, we crossed mice to generate *Chat*^*Cre*^::*Pcdhg*^*1R1/1R1*^::*C4-GFP* animals and analyzed SAC morphology at the individual neuron level (**Figure 3G-K**) and at the population level (**Figure 3B-F**). We found partial rescue by some measurements. By the asymmetry index, SACs expressing the transgene were highly variable, so while they were not significantly better than those from *Pcdhg*^*1R1/1R1*^ mutants without the transgene, they were also not statistically significantly worse than controls (**Figure Figure 3J**). By Elo score however, they were clearly distinct from *Pcdhg*^*+/+*^ but did not separate from transgene-negative *Pcdhg*^*1R1/1R1*^ SACs (**Figure 3K**). Similarly, *Chat*^*Cre*^::*Pcdhg*^*1R1/1R1*^::*C4-GFP* plexus images were not separable from *Pcdhg*^*1R1/1R1*^ by Elo score, but did have intermediate values for interdendritic areas (**Figure 3E-F**).

**Figure 3.**
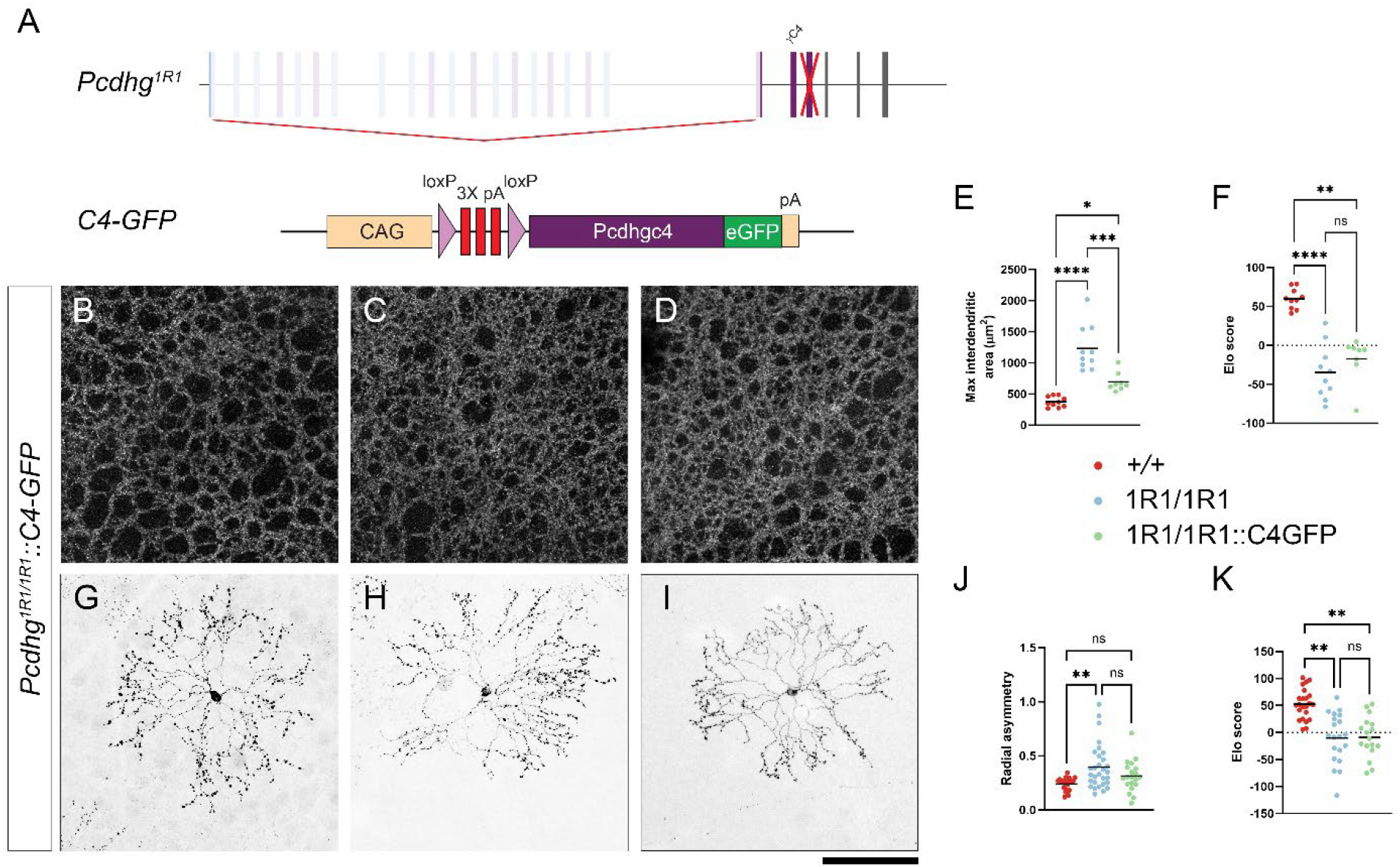
Transgene expression from *C4-GFP* partially rescues self-avoidance in *Pcdhg*^*1R1/1R1*^. **A**) Schematic representation of the *C4-GFP* allele knocked in at the Rosa locus. Expression from the transgene occurs after Cre-mediated recombination of a stop cassette. **B-D**) ON SAC plexus images from *Pcdhg*^*1R1/1R1*^::*C4-GFP* retinas demonstrates interdendritic gaps and dendritic fasciculation intermediate between wild type controls and *Pcdhg*^*1R1/1R1*^ mutants, as quantified in **E** and **F. G-I**) Individual SACs labeled with Brainbow in *Chat*^*Cre*^ also showed incomplete rescue after transgene expression. Quantified in **J-K**. Scale bar is 100 μm. **** is P < 0.0001, *** is P < 0.001, ** is P < 0.01, * is P < 0.05, ns is not significant by Tukey (**E,J**) or Dunn’s (**F,K**) multiple comparison test.

### Normal cell spacing in Pcdhg^1R1/1R1^ mutant retinas

The self-avoidance failures in *Pcdhg*^*1R1/1R1*^ SACs suggest this line could be a useful tool to answer other questions about the role of γ-Pcdhs in directing how neurons arrange themselves during the development. Self-avoidance analyses in the retina have focused on SACs because robust morphological phenotypes have not been described in other retinal neurons in *Pcdhg* mutants. In the retina, neuronal apoptosis and mosaic cell spacing are closely linked, where an increase or decrease in neuronal density may drive errors in spacing and the random distribution of cells throughout the retina. For example, increasing cell density by inhibiting cell death drives some homotypic cells to cluster with each other [35, 36]. Disruption of the entire *Pcdhg* cluster or *Pcdhgc4* alone results in massive apoptosis of amacrine and RGC types via a Bax mediated mechanism [16, 37]. This cell death has made it difficult to test if mosaic spacing relies on γ-Pcdhs as it does on other self-avoidance molecules like the Dscams [38, 39], as the reduced density may mask self-avoidance failures in non-SAC cell types. As the *Pcdhg*^*1R1/1R1*^ mutants have normal cell density [37] but disrupted SAC self-avoidance, we sought to test if previously unappreciated self-avoidance errors were more clear in this mutant.

First, we analyzed the dendrites of vGlut3+ ACs with confocal imaging. These cells densely project to the sublamina between the ON and OFF SACs and undergo excessive apoptosis in whole cluster *Pcdhg* mutants [37]. Investigating either confocal projections through the entire plexus or individual confocal planes (**Figure S1A**) we did not observe any difference between control and *Pcdhg*^*1R1/1R1*^ retinas. Second, to test if mosaic spacing relied on γ-Pcdhs, we analyzed the distributions of dopaminergic ACs, SACs, Vglut3+ ACs, ipRGCs, and Brn3a+ RGCs in control, and *Pcdhg*^*1R1/1R1*^ mutants. We used three metrics to assess spacing and mosaicism for each neuronal type: Nearest neighbor index (NNI), variance of Voronoi domains, and the density recovery profile (DRP), as we have previously used to analyze spacing defects in mutants in the *Dscam* family [34]. For all cell types we did not find any evidence of clustering or loss of mosaic spacing in the absence of 21 γ-Pcdh isoforms (**Figure S1**). In fact, two cell types – ON SACs and Brn3a+ RGCs – became slightly more regularly spaced by the Voronoi measure.

### C4-GFP transgene expression cannot rescue self-avoidance in retinas with both Pcdha and Pcdhg clusters disrupted

Simultaneous disruption of the *Pcdha* and *Pcdhg* clusters caused more severe retinal disorganization than what was observed in mutation of either cluster alone [28]. We hypothesized that targeting α-Pcdhs in our *Pcdhg*^*1R1/1R1*^ mutants would decrease the variability of the self-avoidance defects by making them consistently more severe. To knock down *Pcdha*, we used shRNA sequence previously validated for efficacy and specificity [40]. To verify specificity in our hands, we made a new vector by inserting this shRNA sequence driven by the human U6 promoter as a cassette into a vector encoding eYFP with expression driven by a EF1α promoter and produced AAV particles. We performed RNAscope for *Pcdha* and *Pcdhg* constant exons on retinas transduced with this vector and with a scramble shRNA control. Pcdha was consistently knocked down in YFP-positive cells while Pcdhg was unchanged (**Figure S2**). To specifically analyze SACs, we subcloned this U6-shRNA cassette into Brainbow vectors in the antisense direction to the expression frame to avoid interference (**Figure 4A**). With this organization, the shRNA was expressed in any cell infected by the virus, but the fluorescent reporter was only expressed after Cre-mediate recombination. We injected this vector into the vitreous of P1 pups with three different genotypes: *Chat*^*Cre*^::*Pcdhg*^*+/+*^, and *Chat*^*Cre*^::*Pcdhg*^*1R1/1R1*^, and *Chat*^*Cre*^::*Pcdhg*^*1R1/1R1*^::*C4-GFP*. Knock down of *Pcdha* had no discernable effect on SAC morphology in *Chat*^*Cre*^ mice, but resulted in severe self-avoidance defects in *Chat*^*Cre*^::*Pcdhg*^*1R1/1R1*^ retinas (**Figure 4B-G**). The addition of the C4-GFP transgene did not improve morphologies (**Figure 4H-L**).

**Figure 4.**
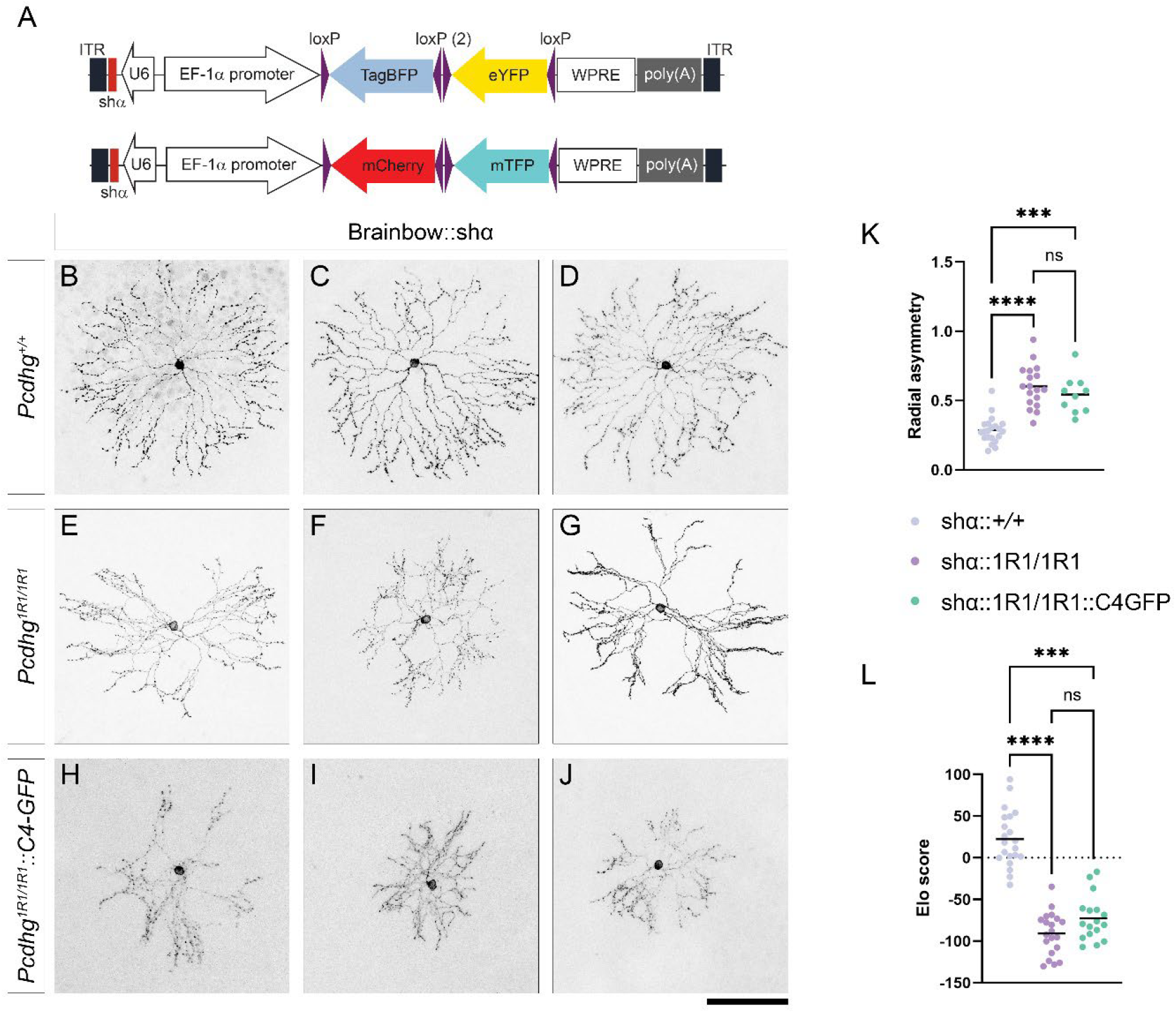
Transgene expression from *C4-GFP* is unable to rescue self-avoidance in SACs with both *Pcdhg* and *Pcdha* disrupted. **A**) Brainbow vectors were modified to include a cassette for shRNA knockdown of *Pcdha* by introducing a human U6 promoter and hairpin RNA sequence antisense to the EF-1α promoter regulating fluorescent protein expression. shα::Brainbow AAV vectors were injected into (**B-D**) *Pcdhg*^*+/+*^, (**E-G**) *Pcdhg*^*1R1/1R1*^, and (**H-J**) *Pcdhg*^*1R1/1R1*^::*C4-GFP* retinas. Self-avoidance defects quantified in **K** and **L** were severe, and the presence of the *C4-GFP* transgene was unable to provide significant rescue. Scale bar is 100 μm. **** is P < 0.0001, *** is P < 0.001, ns is not significant by Tukey (**K**) or Dunn’s (**L**) multiple comparison test.

### γC4-GFP localizes to dendrites and cell-cell junctions in Chat^Cre^::Pcdhg^1R1/1R1^::C4-GFP SACs

One possible reason for incomplete rescue in *Chat*^*Cre*^::*Pcdhg*^*1R1/1R1*^::*C4-GFP* SACs could result from the fact that γC4 does not form homodimers in *cis* but requires a carrier isoform among the other γ-or β-Pcdhs to efficiently localize to the cell surface [5, 7]. In *Pcdhg*^*1R1/1R1*^ with the 21 other γ-Pcdh isoforms absent, there may not be enough carrier isoform available to get γC4 to the surface. To test this hypothesis, we analyzed the localization of γC4-GFP in *Chat*^*Cre*^::*Pcdhg*^*1R1/+*^::*C4-GFP* and *Chat*^*Cre*^::*Pcdhg*^*1R1/1R1*^::*C4-GFP* SACs. We reasoned that we could see more GFP-tagged protein localized to the soma rather than to the dendrites in homozygous mutants if other γ-Pcdh isoforms were required for proper γC4 localization. We analyzed retinas at P14, as self-avoidance is still actively occurring at this time point [33]. We did not discern any differences in localization (**Figure 5A-E**). Expression is high in these transgenic animals and localization to dendrites may not necessarily mean the protein is efficiently inserting into the membrane. However, we also saw pairs of SAC somas adjacent to each other with increased GFP signal at the junctions, consistent with surface localization (**Figure 5F**). These results suggest that γC4 can localize properly in *Pcdhg*^*1R1/1R1*^mutants.

**Figure 5.**
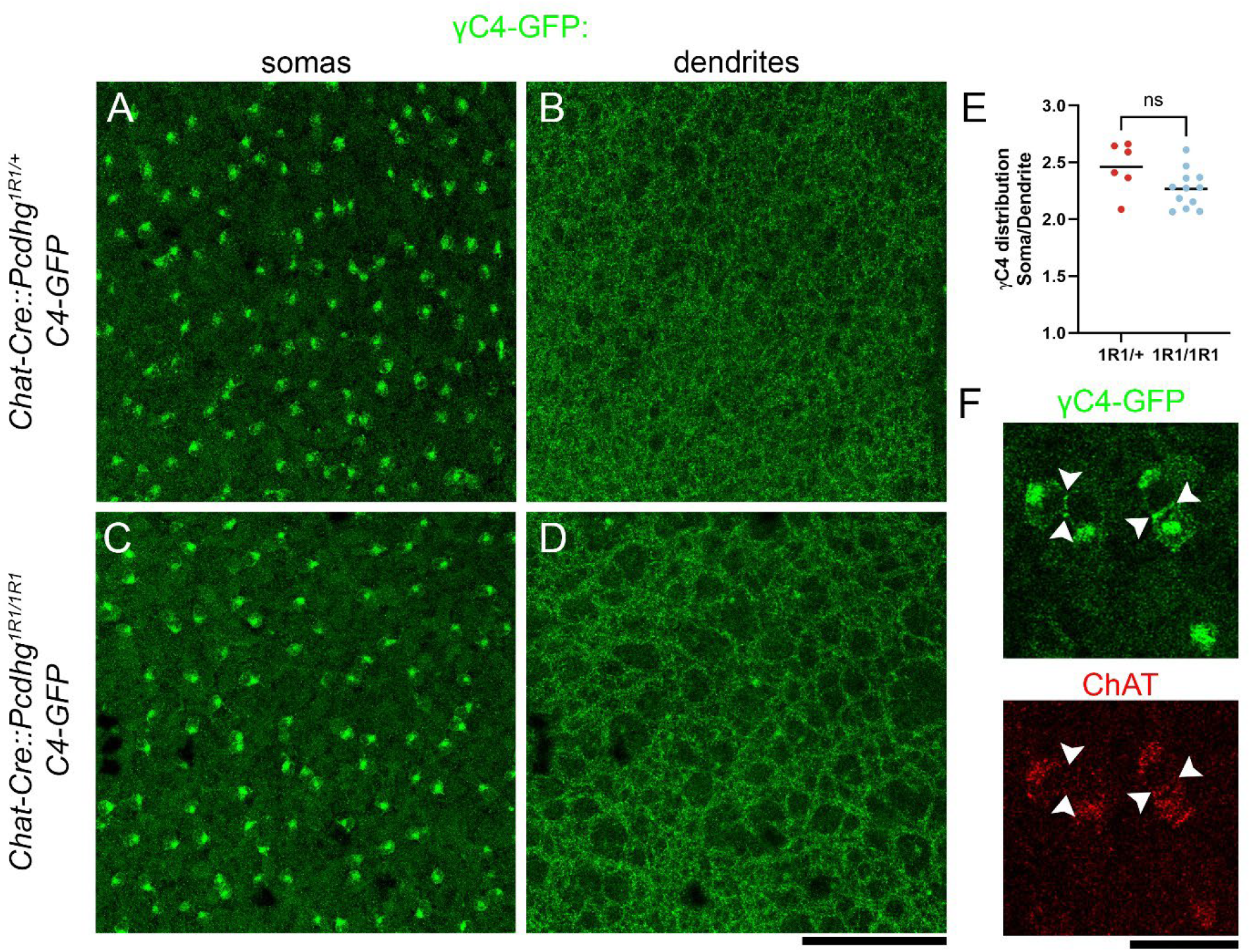
Localization of γC4-GFP does not depend on other γ-Pcdh isoforms. The distribution of GFP-tagged protein in ON SACs was compared between *Pcdhg*^*1R1/+*^ (with one copy of the other 21 Pcdhg isoforms, **A-B**) and *Pcdhg*^*1R1/1R1*^mutants (without any of the other 21 Pcdhg isoforms, **C-D, F**). The ratio of fluorescent signal between somas and dendrites was measured in **E**, with no significant difference between the two genotypes. When SAC somas were adjacent to each other (**F**) there was increased GFP labeling at cell-cell junctions, consistent with γC4 successfully trafficking to the cell membrane and engaging in *trans* interactions in the absence of other γ-Pcdh isoforms.

## DISCUSSION

Here we find that the γC4 isoform of the *Pcdhg* cluster is insufficient to promote neuronal self-avoidance in ON starburst amacrine cells. Previous studies had found that no single isoform was required for SAC self-avoidance [27, 29]. Consistent with that, we saw no defects in SAC morphology in mutants lacking γC4 or γC5. The most parsimonious explanation for this redundancy is that self-avoidance is regulated by a signaling response common to all isoforms mediated by the γ-Pcdh constant domain. Conversely, we found that mice with only γC4 (*Pcdhg*^*1R1/1R1*^) were unable to achieve normal SAC self-avoidance, even with overexpression of γC4 from a transgene. One might have expected similar results to what was found when γC3 or γA1 was overexpressed from a similarly organized transgene: normal individual self-avoidance with reduced non-self contacts [27] and reduced synaptic connectivity between neighboring SACs [29]. Rather, our results were more reminiscent of *Pcdhg* whole cluster loss of function mutants.

We previously found that γC4 is necessary and sufficient to promote neuronal survival [22, 23]. Further, we found that the variable cytoplasmic domain (VCD) was able to rescue survival in an *in vitro* system [37]. Our finding here that γC4 cannot fully promote self-avoidance supports the hypothesis that it has become specialized for that survival function independent of extracellular interactions. γC4 is unique among γ-Pcdhs in that it cannot engage in *cis* homodimers. Goodman and colleagues found that cPcdh *cis* dimers formed asymmetrically with EC6 from one molecule interfacing with EC5/EC6 of the dimer pair. γC4 could only participate as the EC5/EC6 side of the *cis* interaction [7]. This could still allow γC4 to participate in recognition multimers required for self-avoidance. In *Pcdhg*^*1R1/1R1*^mutants with all other γ-Pcdh isoforms absent, γC4 would be primarily dependent on β-Pcdhs for heterodimer partners. β-Pcdhs are intact in these mutants, and we previously reported a compensatory increase in *Pcdhb* isoform expression at the mRNA level in *Pcdhg*^*1R1/1R1*^ animals [22]. Here, we were able to observe γC4-GFP localization consistent with the cell membrane and *trans* interactions between SAC cell bodies (**Figure 5**), but we did not compare its localization with other isoforms. A recent study used γB4 as an example to find that the intracellular domain, including the variable region, can regulate the structure of transmembrane lattices [41]. Thus, we postulate that γC4 cannot efficiently promote self-avoidance because of isoform-specific protein interactions mediated by its VCD, with a potential contribution of unique subcellular localization.

One interesting aspect of our findings is the high variability between individual SACs in the severity of self-avoidance defects. Because of the neonatal lethality in whole cluster mutants, previous studies relied on *Chat*^*Cre*^ to both disrupt *Pcdhg* constant exons with *Pcdhg*^*fcon3*^ and achieve individual SAC labeling with Brainbow [27, 29]. Cre expression from *Chat*^*Cre*^ is sporadic in neonatal SACs before uniformly labeling SACs by ∼P10 [42, 43], but SAC self-avoidance begins earlier than this, and significant differences between wild type and Pcdhg mutants have been observed by P6 [33]. This raised the possibility that variability in SAC self-avoidance could be secondary to variability in when Cre-mediate recombination occurred in individual cells. Since disruption in *Pcdhg*^*1R1/1R1*^ mutants was not dependent on Cre expression, our findings suggest that a substantial amount of this variability resulted from biological causes other than the timing of Cre expression. This does raise an important caveat to keep in mind when comparing the transgene results presented here with those from previous studies. Our “loss of function” mutants were homozygous *Pcdhg*^*1R1/1R1*^animals, while previous studies used Cre to both induce loss of 22 isoforms and introduce single isoforms. This means that, unlike those previous studies, we had mutant cells that never saw transgene expression (i.e., non-cholinergic cells) and mutant SACs that expressed the transgene sporadically before P10.

If γC4 is specialized for neuronal survival to the exclusion of other functions, our *Pcdhg*^*1R1/1R1*^ mutants could be useful for identifying other cell types that require cPcdhs for self-avoidance or other functions for the γ-Pcdhs that were previously masked by the widespread neuronal apoptosis in whole cluster mutants. We began by looking here at cell spacing in several retinal cell types, since spacing is sensitive to changes in cell number [35]. We did not find any major disruption in spacing such as cell clustering as is observed in retinas mutant for other self-avoidance molecules like *Dscam* (**Figure S1**). We also investigated dendrites of Vglut3+ amacrine cells, as their coverage is also influenced by cell number [36], but we did not observe any gross changes. We were also able to increase the severity of self-avoidance failures by knocking down *Pcdha* in *Pcdhg*^*1R1/1R1*^ mutants, which could be a fruitful strategy for affecting multi-cluster disruption in future studies. ON SACs are particularly prone to self-avoidance defects, but future analyses may reveal additional cell types dependent on γ-Pcdhs for self-avoidance or other functions.

## MATERIALS AND METHODS

### Animals

All procedures using animals were performed in accordance with The Guide for the Care and Use of Laboratory Animals and were reviewed and approved by the Institutional Animal Care and Use Committee of Wayne State University. All experiments included a mix of male and female animals. Previously described mouse strains include *Pcdhg*^*1R1*^ [22], *Pcdhg*^*C4KO*^ [22], *Pcdhg*^*C5KO*^ [30], *Pcdhg*^*fcon3*^ (Jax strain number 012644 [14, 16]), *Chat*^*Cre*^ (Jax strain number 031661 [31]), and *C4-GFP* [23].

### RNAscope

Retinas from adult wild type mice were fixed and processed for RNAscope according to the manufacturer’s instructions with the RNAscope Multiplex Fluorescent Assay v2 for fixed-frozen tissue. Probes included *Pcdghc5* (Advanced Cell Diagnostics catalog number 850581); *Chat* (ACD catalog number 408731-C3); *Pcdha* (ACD catalog number 572201-C2); and *Pcdhg* (ACD catalog number 548661).

### Viral vectors

Brainbow viral particles were purchased from Addgene (Catalog numbers 45185-AAV9 and 45186-AAV9). For Brainbow with shRNA knockdown of *Pcdha*, we used the following targeting sequence: AACAGTATCCAGTGCAACACC. We first cloned a U6 promoter into a NotI site with NEBuilder HiFi DNA assembly using a synthesized fragment with the following sequence: CTCCATCACTAGGGGTTCCTGCGGCCGCGAATTCATGATGATGGGATCCCGCGTCCTTTCCACAAGATATATAAACCCAAGAAATCGAAATACTTTCAAGTTACGGTAAGCATATGATAGTCCATTTTAAAACATAATTTTAAAACTGCAAACTACCCAAGAAATTATTACTTTCTACGTCACGTATTTTGTACTAATATCTTTGTGTTTACAGTCAAATTAATTCTAATTATCTCTCTAACAGCCTTGTATCGTATATGCAAATATGAAGGAATCATGGGAAATAGGCCCTCGCGGCCGCACGCGTAAGCTTTGCAAAGATGGA. We then annealed two oligos (AATTTTTCCAAAAAA<u>AACAGTATCCAGTGCAACACC</u>TCTCTTGAA<u>GGTGTTGCACTGGATACTGTT</u>G and GATCC<u>AACAGTATCCAGTGCAACACC</u>TTCAAGAGA<u>GGTGTTGCACTGGATACTGTT</u>TTTTTTGGAAA) and ligated them into a site made by EcoRI/BamHI double digest. AAV particles were generated using the AAV-DJ Helper Free system (Cell Biolabs) as previously described [44]. Briefly, Brainbow-shPcdha constructs were co-transfected with pAAV-DJ and pHelper into 293FT cells (Invitrogen). Transfected cells were lysed three days after transfection, and AAVs were purified with 1-mL HiTrap heparin columns (GE Healthcare).

### Intravitreal injection

For Brainbow labeling, adult mice were anesthetized by isoflurane. A sharp 27G needle was used to make hole in the sclera below the limbus and 10^9^ vg AAV particles in 1μl was injected into the vitreous using a blunt 33G Hamilton needle. AAV vectors for shRNA knockdown were introduced at P1. Pups were anesthetized on ice, a sharp needle was used to open the eyelid and puncture the sclera, then a blunt Hamilton needle was used to introduce 0.5μl AAV into the vitreous. At both ages, topical proparacaine was used for analgesia after injection. Retinas were collected for analysis 2-3 weeks after injection.

### Immunohistochemistry

Eye cups were fixed by immersion in 4% PFA for 1-3 hours with cornea and lens removed, then rinsed in PBS. Retinas were dissected in PBS. Primary antibody staining was performed for 48 hours at 4°C in blocking solution containing 2.5% bovine serum albumin (BSA) and 0.25% Triton-x-100 in PBS, and secondary antibody staining was performed in the same solution overnight. The following antibodies were used at the indicated concentrations: Goat anti-ChAT, 1:500 (Sigma-Aldrich Cat# AB144P, RRID:AB_2079751); Chicken anti-GFP, 1:2000 (Aves Labs Cat# GFP-1010, RRID:AB_2307313); Rabbit anti-mCherry, 1:1000 (Millipore Cat# AB356482, RRID:AB_2889995); Alexa Fluor 647 conjugated Alpaca anti-TagFP, 1:500 (NanoTag Biotechnologies Cat# N0502-AF647-L, RRID:AB_3075936); Guinea Pig anti-Vglut3, 1:10,000 (Millipore Cat# AB5421, RRID:AB_2187832); Mouse anti-Brn3a, 1:500 (Millipore Cat# MAB1585, RRID:AB_94166); Rabbit anti-melanopsin, 1:2000 (Advanced Targeting Systems Cat# UF006, RRID:AB_2314781). Secondary antibodies were Alexa Fluor conjugated and used at 1:500.

### Image analysis and statistical comparison

Whole mount retinas were imaged *en face* with a Lecia Sp8 confocal microscope. For individual SACs, z-projections were compiled in Fiji and their radial symmetry measured using the Azimuthal Average plugin. This method measures the variance in fluorescent signal between eight radial bins centered on the SAC soma. Values across genotypes were compared by one-way ANOVA with Tukey’s post hoc comparisons.

For Elo analysis, we used the ImageEchelon tool available here: https://github.com/TheJacksonLaboratory/ImageEchelon and described in detail previously [45]. This tool simply selects random image pairs from a dataset and presents them to a trained observer who chooses which of the two is the “winner” according to whatever parameter is set by the experiment. In this case, the SAC with the more intact self-avoidance was deemed “winner”. After the observer selects the winner, ImageEchelon updates image scores based on an Elo algorithm such that the “winner” gains points equal to the value lost by the “loser” image. More points are exchanged if the “winner” entered the matchup with the lower starting score (i.e., as the underdog). We have found that this method can efficiently rank images without performing every pairwise comparison. We continued comparisons until the average matchup number was 25. For individual SACs, each neuron was considered a biological replicate. All conditions were compared in a single set of images. Since Elo scores are ordinal, we compared between genotypes using a Kruskal-Wallis test with Dunn’s multiple comparisons. Summary statistics for individual SACs with all pairwise comparisons are in supplementary file 1. For Elo comparisons of ON SAC plexus images, observers were instructed to choose images with more diffuse dendrite coverage rather than those with fascicles and gaps. For these, 2-4 images were scored per retina (technical replicates). Scores were averaged to yield a single score per retina (biological replicates). Retinas were compared across genotypes using Kruskal-Wallis with Dunn’s posthoc test for multiple comparisons. Summary statistics with all pairwise comparisons are in supplementary file 1.

Maximum interdendritic area was measured in Fiji by sampling multiple candidate gaps within an image to identify the largest. Values from 2-4 images per retina were averaged to yield a single number per retina. These values were compared between all genotypes by one-way ANOVA with Tukey’s post-hoc test.

To analyze the distribution of γC4-GFP in SACs, we stained for GFP in P14 retinas expressing the transgene in cholinergic cells. We collected confocal stacks spanning from the RGL through the IPL and made sub-stacks with 10 steps through the somas and 5 steps through the ON plexus.

Each sub-stack was normalized identically using the “Subtract Background” command in Fiji with a rolling ball radius of 50 pixels. In the soma sub-stack, we then used a circular ROI with 15.9 μm diameter to measure the mean fluorescence intensity of all 10 steps centered on a soma. These values were summed to approximate the fluorescence intensity through a single soma. Ten somas per image were measured and their values averaged to yield a single number per image. For the plexus sub-stack, the mean intensity through the entire field was summed over the 5 steps to approximate the fluorescent signal with the dendrites. The ratio of these values (soma/dendrite) was used to approximate the relative distribution of the protein and compared between genotypes with a t-test.

Statistical analyses above were performed in GraphPad Prism. Outputs from all statistical tests are in Supplementary File 1.

Cell spacing was measured by first defining x,y coordinates for each cell body using the cell counter plugin in Fiji. An R script was used to quantify spacing from these coordinates by three methods: the sjedrp calculated Density Recovery Profile (DRP) based on previous methods [46]; Voronoi domains were calculated using the deldir package according to previous methods [47], then the variance of the area of the domains was calculated as Voronoi score; Nearest neighbor regularity index (NNRI) was calculated using the SpatialEco package. Both Voronoi and NNRI values were compared between wild type and *Pcdhg*^*1R1/1R1*^ using a t-test.

## ACKNOWLEDGEMENTS

This work was funded by NIH/NEI R01 EY031690 and R21 EY033874 to A.M.G. and by Career Start Grant from the Knights Templar Eye Foundation to S.S. The WSU Vision Research Core is funded by P30 EY004068 (Core Grant to Linda Hazlett, Ph.D.) and an unrestricted grant from Research to Prevent Blindness to the Department of Ophthalmology, Visual and Anatomical Sciences at WSU SOM. We thank Bikash Rana and Stephanie Garrett for assistance with image analysis, and Dr. Joongkyu Park for assistance with AAV production technique.

