## Supplemental Figures for "Reduced neuronal self-avoidance in mouse starburst amacrine cells with only one *Pcdhg* isoform"

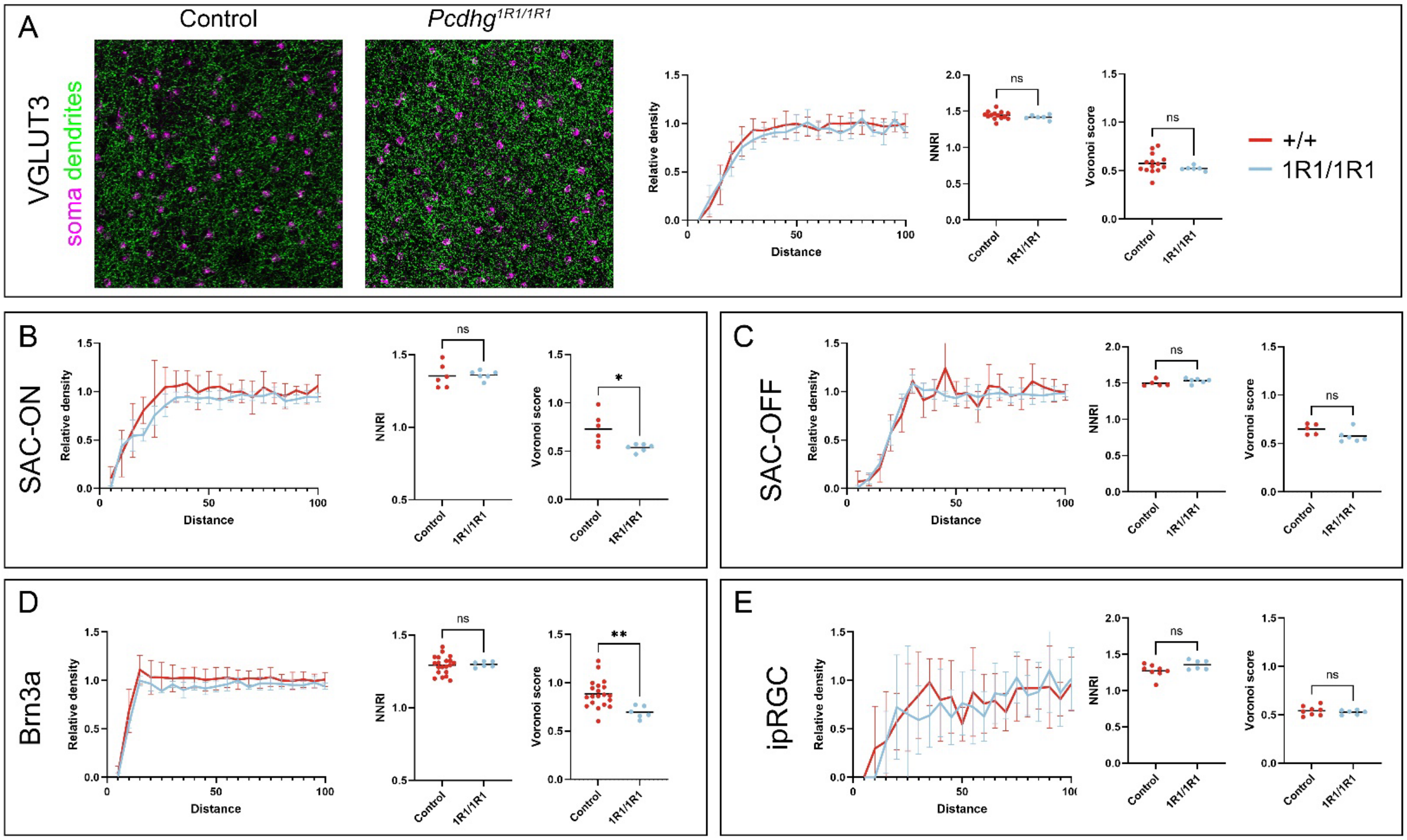


**Figure S1: Normal cell spacing in *Pcdhg*$^{1R1/1R1}$ retinas.** To ask if other cell types showed phenotypes reminiscent of self-avoidance failures with normal cell number in *Pcdhg*$^{1R1/1R1}$, we analyzed dendrite coverage and cell spacing in (**A**) Vglut3+ amacrine cells as well as cell spacing in (**B**) ON SACs, (**C**) OFF-SACs, (**D**) Brn3a+ RGCs and (**E**) melanopsin+ ipRGCs. No cell type showed abnormal clustering or loss of mosaic spacing in *Pcdhg*$^{1R1/1R1}$.

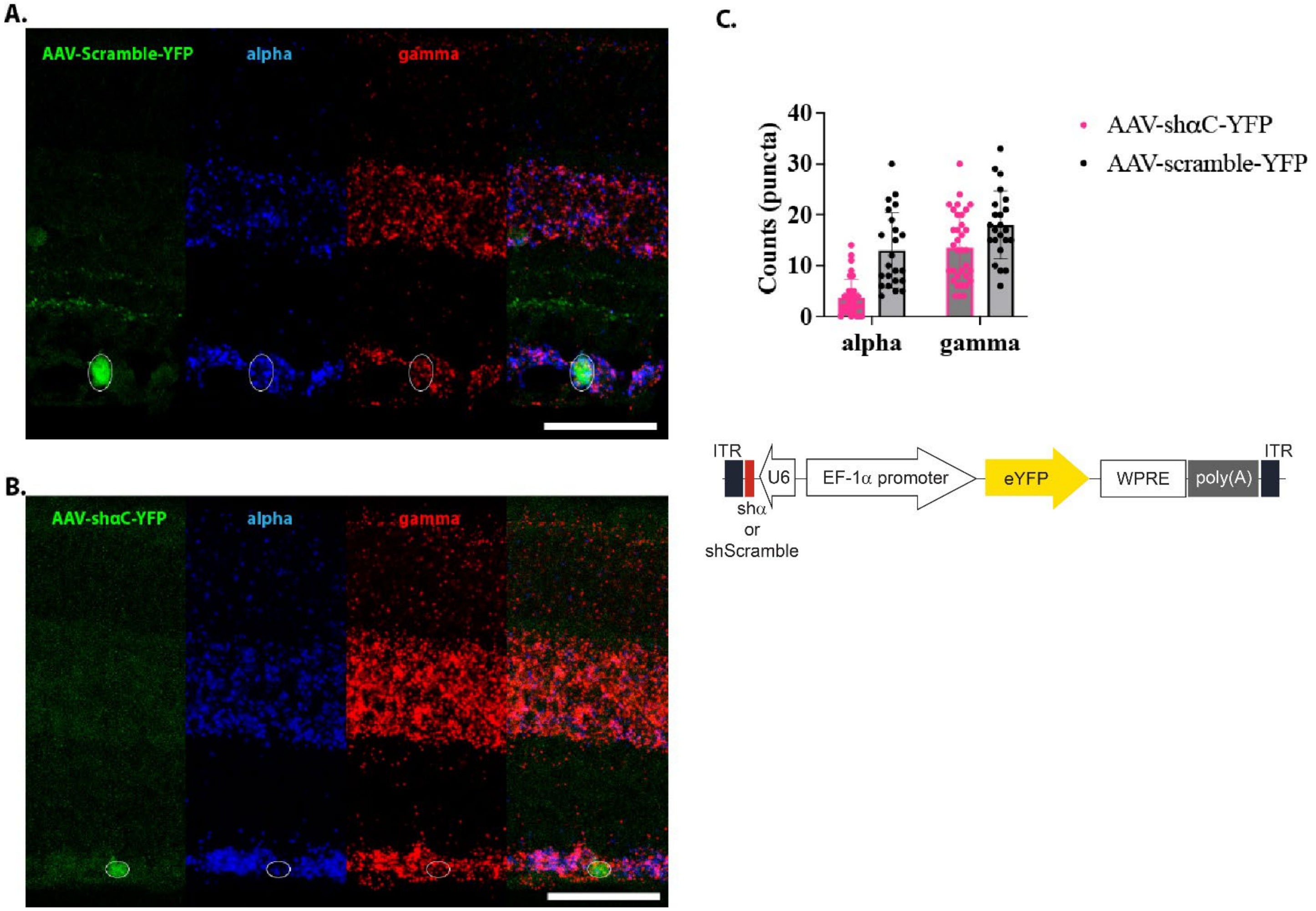


**Figure S2: Verification of *Pcdha* knockdown.** Retinas were injected with (**A**) AAV-shScramble or (**B**) AAV-shαC and processed for RNAscope using probes against *Pcdha* constant domain (blue) or *Pcdhg* constant domain (red) along with antibody co-detection for YFP. ISH puncta within YFP-positive cells were counted as an estimate of expression and quantified in **C**. Schematic of the shRNA construct is in **D**. Scale bar is 50 μm.
